# Conclusions Drawn From Neural Network to Brain Alignment Depend Strongly on the Chosen Similarity Measure

**DOI:** 10.1101/2024.08.07.607035

**Authors:** Ansh Soni, Sudhanshu Srivastava, Marvin Maechler, Konrad Kording, Meenakshi Khosla

## Abstract

Deep neural networks are widely used to model biological perception and behavior, making the similarities and differences between artificial and biological systems consequential. If a principle (e.g. self-supervised learning) produces a model resembling biology, this is taken as evidence the same principle shapes the biological system. But what it means for an artificial system to be similar to a biological one is complex. A popular approach compares representations of identical stimuli using a similarity measure like Representational Similarity Analysis. Yet scientific questions rarely specify which measure is appropriate, raising a key question: do conclusions depend on this choice? Focusing on vision, we show that measure choice influences both hierarchical correspondence between systems and the ranking of which artificial systems are most biological. Reanalyzing prior studies, we find the choice hugely consequential: models best under one measure are often worst under another, and prior conclusions can flip or dissolve. Different metrics capture fundamentally different aspects of similarity, warranting healthy skepticism toward such comparisons.

## 1 Introduction

Researchers have built artificial neural models of the brain since McCulloch and Pitts debuted their model of a neuron in 1943 [1]. In the last decade, deep neural networks (DNNs), our most complex artificial neural systems, have allowed researchers to make computational models that can not only model internal mechanisms, but come close to matching some aspects of the perception and behaviour of biological neural systems [2–6]. In vision, the focus of this paper, DNNs have been the artificial neural system of choice to model a wide range of perceptual and behavioral phenomena such as eye movements [7], category selectivity [8–10], and complex responses on visual tasks [9, 11]. A central observation that motivated this research program is that artificial neural systems trained on object categorization develop internal representations that resemble those in the primate ventral visual stream [2, 12], leading to the proposal that brains and artificial networks may share common computational goals [13–15].

The method of comparing internal representations solved a fundamental problem in evaluating these artificial neural systems, representing a new way to deduce general scientific insights about biological neural systems. Since researchers found a correspondence between the internal representations of biological and artificial neural systems, the inductive biases that shape the artificial neural systems (i.e. architecture, learning rule, dataset) should determine the exact level of this correspondence. Therefore when the choices we make in shaping the artificial neural system increase their correspondence to biological ones, we conclude that these inductive biases have also shaped the biological ones. To measure this correspondence, researchers present the same stimuli to a biological neural system (i.e. a human subject) and to multiple artificial systems each using different methods to encode candidate inductive biases that may shape the biological neural systems (i.e. instance- versus category-level training), the internal representations of these neural systems are extracted, and then a *similarity measure* is computed, providing a scalar that quantifies the correspondence of these representations (Fig. 1).

**Fig. 1.**
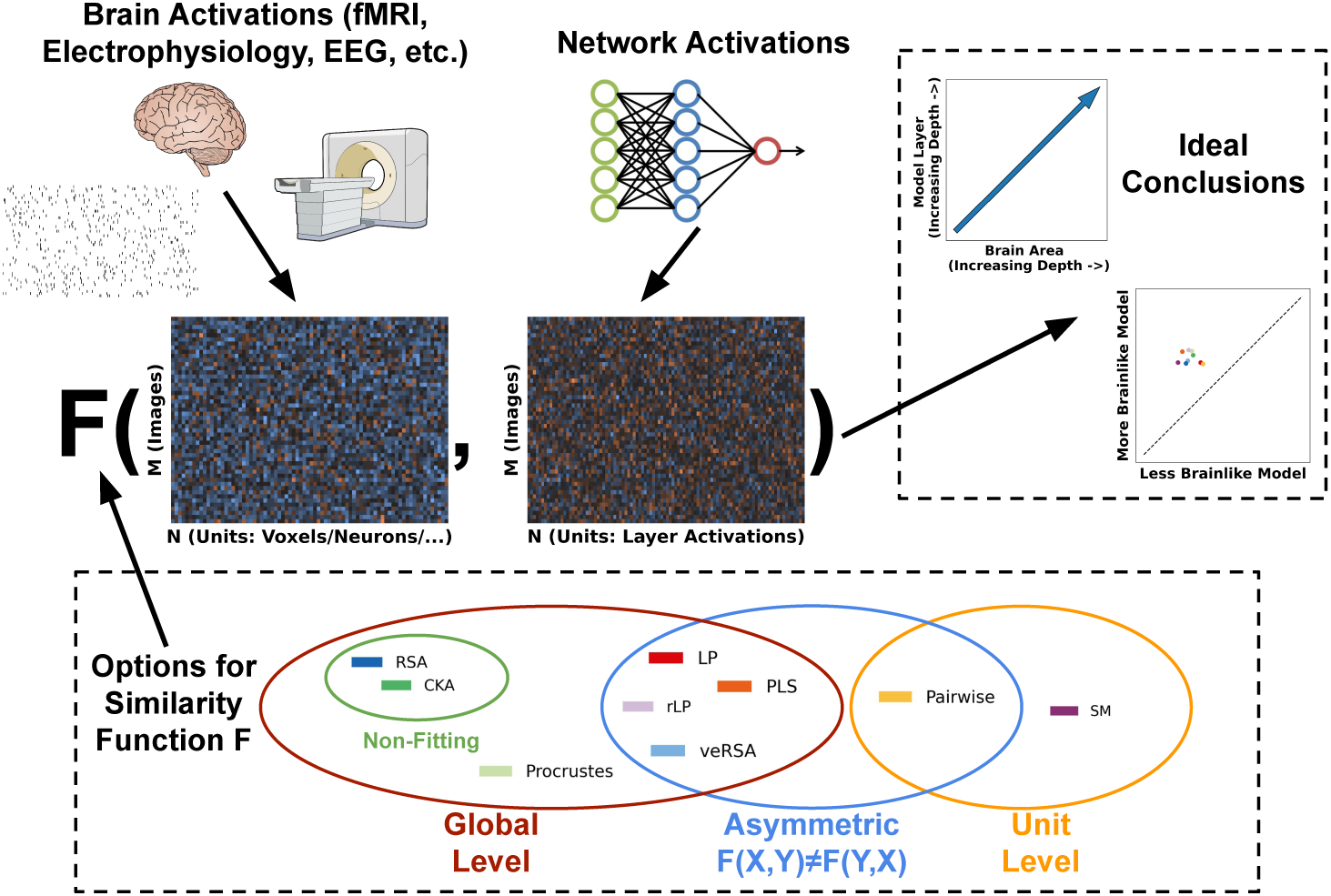
Asking how the choice of measure matters when comparing brain data with neural network activities. Activations from Brain and Model are extracted for a shared N Stimuli utilizing various methods (e.g., single-unit recordings, MEG, EEG, fMRI). These activations are compared using a function that outputs a similarity score. Some choices for this function are listed (9 chosen for this paper). Measures have various theoretical similarities, and the main differences are highlighted. These similarity scores are then used to make various conclusions, such as which network is better and hierarchical correspondence. An ideal example is shown. Images used are sourced from commons.wikimedia.org

Building on these comparisons, the field has grown into a substantial research program. Large-scale neural datasets [16–18], public benchmarks [3, 19–21], have allowed a steady stream of papers to test how specific design choices for artificial neural systems — supervision [22], language training [23, 24], adversarial robustness [25–27], and spatial regularization [28] — affect their correspondence with biological ones, using these results to make claims about the inductive biases that shape these neural systems. These comparisons are the primary empirical basis for claims about which computational principles make a network more *brain-like*.

A fundamental assumption for this method to make principled scientific conclusions is that the score provided by the similarity measure is able to capture the effect of the choices made that are linked to a specific inductive bias. This, arguably, implies either that the specific choice of this measure is the only theoretically justifiable one, or that any theoretically justified measure will lead to similar scientific conclusions (up to some level of noise). While some papers directly link their choice of measure to their hypothesized inductive biases (e.g. [9]) the majority provide a justification that does not make a specific link, instead providing a principle that would allow many measures to be equally justifiable. For example, some papers set their scope to rule out measures that are too permissive (e.g. [28]) or provide a general property on how units should be mapped between two systems (e.g. [22]). With a large number of different measures that are in use and would fit these properties [29], we should expect that they all give similar results.

However this assumption of the irrelevance of the used metric has yet to be tested. Individual studies almost always report a single measure, and the specific implementation of that measure often varies between papers [30]. The assumption that all justifiable measures will lead to the same conclusions has not been tested systematically. Because each measure has a different theoretical rationale and different mathematical properties, there is no guarantee that they agree. If they do not agree, unless the chosen measure is tightly linked to the question, it is the measure rather than the underlying hypotheses that determine what the literature concludes.

Here we test this directly. We compute nine similarity measures on a common set of models, brain datasets, and pipelines, and ask whether they agree on the questions the field actually tries to answer: which layer of a network best matches which brain area, which of a set of candidate models is most brain-like, and whether training manipulations such as self-supervision or multimodal language training bring models closer to the brain. They do not agree. Different measures produce different layer-to-area mappings, different model rankings, and, when applied to the datasets and models of several prominent prior studies, lead to different qualitative conclusions. This goes well beyond an expected level of noise as we see cases where different measures reach statistically significant results in opposite directions. The choice of measure is not a technicality.

Instead, it is often the dominant factor determining what a model-brain comparison concludes, and claims about the relative brain-alignment of neural networks should not be treated as established findings about the brain. The field needs both a more careful theoretical account of when specific measures are appropriate and a default practice of reporting multiple measures tightly linked with a statement of the *nature* — not only the degree — of any similarity that is observed.

## 2 Methods

We implement 9 different measures. Each measure takes 2 matrices with a shared first dimension, M (stimuli) × N (features) and outputs a similarity score. Non-Fitting measures such as representational similarity analysis (RSA [31]) and centered kernel alignment (CKA [32]) output a single score. Fitting measures like Linear Predictivity and voxel-encoded RSA (veRSA, [22]) output multiple scores as a result of a K-Fold (5-fold) validation which are then averaged for a final score. Comparisons are repeated for every subject within the dataset which are then averaged again. This gives 9 scores, one for each measure, for every dataset-model combination.

We also implement Sparse Random Projection as a possible dimensionality reduction step following the process of [33]. We utilize the Johnson-Lindenstrauss (JL) lemma [34] which takes the number of stimuli and returns the minimum number of dimensions required to preserve the Euclidean distance between points within a factor of 1 ± *ɛ* which we set to *ɛ* = .1, the number of dimensions varies depending on the number of images in the dataset. Comparisons with no dimensionality reduction were also completed.

Model activations were typically extracted after every block instead of every layer for comparison to brain data. For a more thorough comparison, we extracted activations for comparison after every layer for AlexNet.

We chose these datasets/models/metrics on a combination of popularity, availability, and on the basis of the existing results we were looking to reproduce/reanalyze.

### 2.1 Metric Implementation Details

#### 2.1.1 Linear Predictivity (LP)

For Linear Predictivity (LP), we fit a Ridge regression with the model neuron responses as the predictor variables, to predict brain unit responses using 80% of M. The RidgeCV function in Scikit-learn was used with the regularization parameter, *α*, chosen as the optimal parameter from a logarithmic space between 10^−8^ and 10^8^. The similarity score is then calculated as the *R*^2^ on the held-out 20% of *M* unseen images.

LP asks if the brain’s responses can be linearly read out from the features present in the model’s representation. It is best when you care about what is decodable. The main weakness of LP is its leniency. A model with a larger set of rich features may score well without truly matching the brain, and there is no penalty for the model carrying extra information that the brain lacks.

#### 2.1.2 Reverse Linear Predictivity (rLP)

Reverse Linear Predictivity (rLP) was calculated using the same procedure as Linear Predictivity, with the voxel responses being used as the predictor variables to predict model unit responses. Albeit not a commonly used measure, it has seen some use in the past [35].

rLP flips the direction of LP, asking if the model’s features can be linearly read out from the brain. This penalizes models that encode things the brain does not. However as neural data comes from noisy or sparse measurements, a low score can be due to measurement limits instead of a real mismatch.

#### 2.1.3 Partial Least Squares Regression (PLS)

Partial Least Squares regression (PLS) was fit using the PLSRegression function in Scikit-learn with 25 components (chosen to match previous studies [2, 19]), to predict voxel responses using model neuron responses. Similar to the other predictivity metrics, we used a 80/20 split on M images.

PLS is a more cautious version of LP. It only uses the most dominant patterns between model and brain, guarding against overfitting from oversized feature sets. The catch is the fixed number of patterns it keeps is a choice to be made by the researcher. Setting the number too low throws away real signal, and too high makes it as lenient as plain LP.

#### 2.1.4 Representational Similarity Analysis (RSA)

Representational Dissimilarity Matrices (RDM) of size *M* ×*M* were first computed for model responses as well as voxel responses by calculating the pairwise dissimilarity (1 - correlation) for each pair of images. The similarity score was then computed using the Kendall’s *τ* correlation between the upper triangle of the two matrices (Spearman’s *ρ_a_* is an alternative if an analytical noise ceiling is desired, [36]). RSA has many variants as any distance metric can replace 1 - correlation for the RDMs and any similarity metric can replace Kendall’s *τ*. This has led to some formulations that are equivalent to other metrics [37].

The traditional correlation RSA (which we use here) asks if two systems organize stimuli in a similar way, specifically what stimuli are considered alike or different. It requires no fitting, making it simple to utilize. The weakness is that two systems can have matching RDMs even if one encodes a feature the other ignores (e.g. object size or viewpoint) as long as that feature does not change the relative distances between stimuli.

#### 2.1.5 Centered Kernel Alignment (CKA)

We used linear Centered Kernel Alignment (CKA), proposed in [32]. Linear CKA calculates the similarity between two matrices *X* ∈ R*^M^*^×^*^Nx^* and *Y* ∈ R*^M^*^×^*^Ny^* of neuronal responses using: 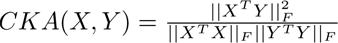 where *F* denotes the Frobenius norm. The matrices *X* and *Y* have centered columns.

CKA is a quick comparison of how two systems organize their representations, needing no fitting. However this quick, general comparison can lead to a few dominant patterns driving the score, so its similarity may be high while ignoring finer structure. Alternatively, it can be thrown off by a small number of unusual stimuli.

#### 2.1.6 veRSA

Voxel Encoded RSA (veRSA) follows the same ridge regression procedure as Linear predictivity. However, instead of computing the *R*^2^, a Representational Dissimilarity Matrix (RDM) is generated from predicted values on the test set, and RSA (with the same process as described above) is then applied to obtain the final score.

veRSA first reweights the model’s features to best match the brain, then compares geometry the way RSA does. This allows a model to map features to the brain even when the brain data is combining across many units (e.g. fMRI). That flexibility is also its weakness, as it can hide real mismatches.

#### 2.1.7 Procrustes Distance

Let *X* ∈ R*^M^*^×^*^Nx^* and *Y* ∈ R*^M^*^×^*^Ny^* be the response matrices. To compute the Procrustes distance, first both *X* and *Y* are mean-centered and normalized. They are then projected onto a common subspace of dimension *N_d_* = min{*N_x_, N_y_, M* } via principal component analysis (PCA), as recommended in [38]. Next, we compute an orthogonal matrix *T* in the orthogonal group

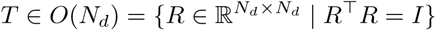

that best aligns *Y* to *X*, obtained in closed form from the singular value decomposition of *X*^⊤^*Y*. That is, we define the Procrustes distance as the angular (geodesic) shape distance between the aligned representations,

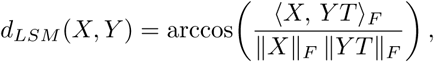

where ⟨*A, B*⟩*_F_* = Σ*_i,j_a_ij_b_ij_* and the Frobenius norm is given by 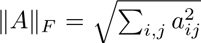 The alignment is carried out using the procedure outlined in [38]. Finally, the distance score *d_LSM_* (*X, Y*) is negated to obtain a similarity measure.

Procrustes is a stricter geometric comparison compared to the previous measures. It tries to match up two representations using a single orthogonal transform, asking if they are the same shape. The trade-off is that strictness, as real correspondences may be scored as dissimilar if they require a more flexible transform, and it requires similar dimensions, or the initial dimensionality-reduction step can discard some structure.

#### 2.1.8 Pairwise matching

Pairwise matching compares two populations *X* ∈ R*^M^*^×^*^Nx^* and *Y* ∈ R*^M^*^×^*^Ny^* by first calculating, for each neuron in *X*, the best matching neuron in *Y* by calculating their response correlations on a training set of stimuli. Next, the correlation of each neuron in *X* with its best-matching neuron in *Y* is calculated on a testing set with the final score as follows:

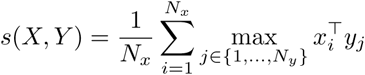

The procedure for calculating the pairwise matching is similar to the implementation presented in Khosla et. al. [9].

Pairwise matching is the strictest measurement, caring about unit-by-unit correspondence. It asks if each model unit has a counterpart in the brain. It suits questions about whether the entire selectivity profile of a unit is preserved. Its limitation is that it privileges a single-neuron perspective so systems with similar population-level geometry can receive lower scores if the same manifold is oriented differently with respect to the neural axes.

#### 2.1.9 Soft Matching (SM)

As opposed to pairwise distance, Soft Matching (SM) is symmetric. Soft Matching is calculated as the following:

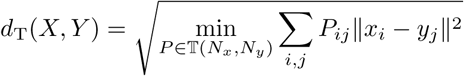

where the transport matrix **P** ∈ R*^Nx^*^×^*^Ny^* is a non-negative matrix that has rows which sum to 1*/N_x_* and columns which sum to 1*/N_y_*. These matrices belong to a *transportation polytope* [39], denoted as T(*N_x_, N_y_*). This implementation is described further in Khosla et. al. [9]. This distance score is made negative to become a similarity. Soft Matching is sensitive to rotations while not caring about how units are labeled.

It asks whether two systems share the same preferred axes, meaning the tuning of individual units, rather than just the overall geometry. This lets it pick up structure that geometry-based metrics miss. It is symmetric and it avoids treating a network with many units as similar to everything. The trade-off is its strictness, since it penalizes models that carry the same information spread across a different set of units.

### 2.2 Error Bars and Reference Points

Error bars are included as ± Standard Error where possible across subjects and/or k-fold validation to follow standard practice. Like most other papers, these error bars ignore multiple sources of error. For example, multiple viewings in NSD are often averaged before this comparison is completed. This and other sources of error are not accounted for in the error bars.

We computed a “maximum” reference point (sometimes referred to in the literature as a noise ceiling) through inter-subject similarity for each measure. This was computed utilizing the same procedures as in model-brain comparisons, instead holding one subject out as the current “model” and computing scores with the remaining subjects, repeating this for each subject being the “model”. This is the same as taking the average score for every pairwise comparison between the subjects, repeated for each metric. This “maximum” also has varying ways of being calculated. For example some papers have used a split-half validation (across subjects), leave-one-out validation (across subjects), or across stimuli repetitions (across trials). This leads to significant differences in interpreting raw scores of models across papers and can over- or under-inflate how close we are to understanding the brain. Differing reference calculations have also caused varying conclusions in the past, with [40] remedying an incorrect calculation with one that is closer to the intersubject version we use, with a bootstrapping step. This new reference caused the work to conclude “The claim that the model “fully explains” the human IT data appears overstated” causing a significant change to the conclusions made by the work. More work is clearly required to standardize these reference points, on top of how to use the metrics to ensure consistent scientific conclusions. A more thorough discussion of this problem is available in [41], which also includes proper methods on computing various reference points for specific metrics.

We also computed two types of “minimum” reference points or alternative baselines for NSD: First using the raw pixel values of the image by vectorizing it, and another utilizing the category information of the images by using a vector of classes with elements corresponding to the number of a class present in the image, this vector is then normalized to sum to 1.

### 2.3 Variations in Metrics Over Many Papers

When choosing the implementations of each metric, we chose hyperparameters that best matched recent papers in the field. We keep this consistent throughout the paper, leading to small differences in measure hyperparameters between the comparisons presented here and the original study implementations. For example, the 3 measures used in Margalit et al., 2024 [28] vary slightly from our implementation. First, their linear regression measure was a PLS regression with 1000 components, while we used 25 components (following previous studies with shared authors [2, 19]). Second, when computing RSA, they collapsed across 5 categories (faces, bodies, characters, places, objects), whereas we used the separate 515 images. Finally, our pairwise matching metric, while similar to their one-to-one metric, is slightly more permissive, allowing the same unit in one representation to be mapped to multiple different units in another representation. If small changes in hyperparameters lead to large differences in comparisons, there should be a much larger focus on why these hyperparameters are selected.

### 2.4 Models Used

#### 2.4.1 AlexNet

AlexNet [42] is a small CNN, one of the first models in deep learning. We used the default PyTorch implementation ([43]) of AlexNet pre-trained on ImageNet.

#### 2.4.2 VGG

VGG [44] is a deeper CNN architecture with multiple depth options. We used the default PyTorch implementation ([43]) of VGG networks pre-trained on ImageNet.

#### 2.4.3 ResNet

ResNet [45] is a deep CNN architecture with residual connections and multiple depth options. We used the default PyTorch implementations ([43]) of ResNet networks pretrained on ImageNet with supervised learning.

#### 2.4.4 Instance-Prototype Contrastive Learning (IPCL)

We used the pre-trained model weights from Konkle and Alvarez, 2022 [22]. This included Alexnet models with category-supervised, self-supervised (IPCL), and randomly initialized versions.

#### 2.4.5 Vision Transformers (ViT)

ViT is a transformer-based network [46]. We used multiple sizes of the default PyTorch implementation ([43]) of ViT trained on ImageNet with category-supervision.

#### 2.4.6 Topographic Deep Artificial Neural Network (TDANN)

We used the pre-trained model weights from Margalit et al., 2024 [28]. This included various ResNet18’s trained with different objectives. These objectives were a combination of self-supervised and category-supervised combined with a spatial loss to incentivize locally correlated units.

### 2.5 Datasets Used

We utilize the processed data for all datasets and therefore omit these details. Detailed descriptions of these steps are available in the cited papers.

#### 2.5.1 Natural Scenes Dataset

Detailed Descriptions of the NSD dataset and its collection can be found here [16]. This dataset contains 7T fMRI data (1.8 mm, 1.6 s) for 8 participants, each viewing 9-10 thousand images multiple times totaling 22-30 thousand trials. Subjects completed a long-term continuous recognition task while viewing the images. The images were taken from the Common objects in context dataset ([47]), a set of natural images created for computer vision tasks. We extracted ROI’s from the data separating out V1,V2,V4, the rest of the Ventral stream (labelled VVS in plots), Dorsal stream (labelled DVS in plots), and Lateral stream (labelled LVS in plots). We used both the shared 515 images with 8 participants and shared 1000 images with 4 participants.

#### 2.5.2 Object Orientation Dataset

Detailed Descriptions for this dataset can be found in Konkle & Alvarez 2022 [22]. This dataset contains 3T fMRI data (1 mm,2.2s) for 4 participants, each viewing 40 images 4 times for a total of 160 trials. The images were of 8 items presented at different orientations from 0 - 180 degrees spaced equally. Subjects completed a vigilance task by pressing a button when a red circle appeared.

#### 2.5.3 Inanimate Objects Dataset

Detailed Descriptions for this dataset can be found in Konkle & Alvarez 2022 [22]. This dataset contains 3T fMRI data (3 mm,2.0s) for 8 participants, each viewing 72 images 6 times for a total of 432 trials. The images were of 72 different inanimate objects. Subjects completed a vigilance task by pressing a button when a red frame appeared.

#### 2.5.4 Monkey V1 Dataset

Detailed Descriptions for this dataset can be found in Cadena et al 2019 [48]. Single neuron recordings to 7250 images for 2 monkeys. This dataset consists of ImageNet images [49] along with texturized versions. Stimuli were repeated 4 times. Due to more repeats than subjects and the low number of neurons (51 and 115), errors were taken across repeats as opposed to subjects.

#### 2.5.5 ManyMonkeys Dataset

Detailed Descriptions for this dataset can be found in Dapello et al 2023 [27]. These datasets contain single neural recordings to 640 images. The images are synthetic “naturalistic” images of single objects. Micro-electrode arrays were placed in the IT cortex of 6 monkeys leading to a varied number of 58-280 sites from each monkey. The firing rate was averaged over a 70-170 ms window following the onset of stimulus presentation.

## 3 Results

### 3.1 The hierarchical mapping between layers and brain areas depends on the measure and does not replicate between brains

A foundational finding in NeuroAI is that task-optimized DNNs replicate the hierarchical organization of the primate visual cortex [50, 51]: successively deeper layers of the network best align with successively higher visual areas. This mapping is one of the main reasons DNNs are taken seriously as mechanistic models of vision, because it suggests that the computations that these artificial neural systems do match biological ones and more specifically that what deep layers compute is in some sense what higher visual areas compute. We find that the basic pattern — early layers best aligning with early visual areas (V1, V2) and later layers best aligning with higher visual areas (IT) — is recovered by many but not all measures, and the specific layer each measure assigns to each area varies (Fig. 2). Hierarchical correspondence breaks down entirely for soft matching, pairwise matching, and reverse linear predictivity, which tend to collapse onto one or two layers rather than showing a progression with cortical depth. This is consistent with recent between-model work, where soft matching performs worse than linear predictivity and RSA at clustering corresponding layers across models with similar architectures and training [52]. Hierarchical correspondence has also been shown to be different for different models (especially ones that include skip connections) [53] and we see similar variance, however the effect of measure seems to dominate in our comparisons.

**Fig. 2.**
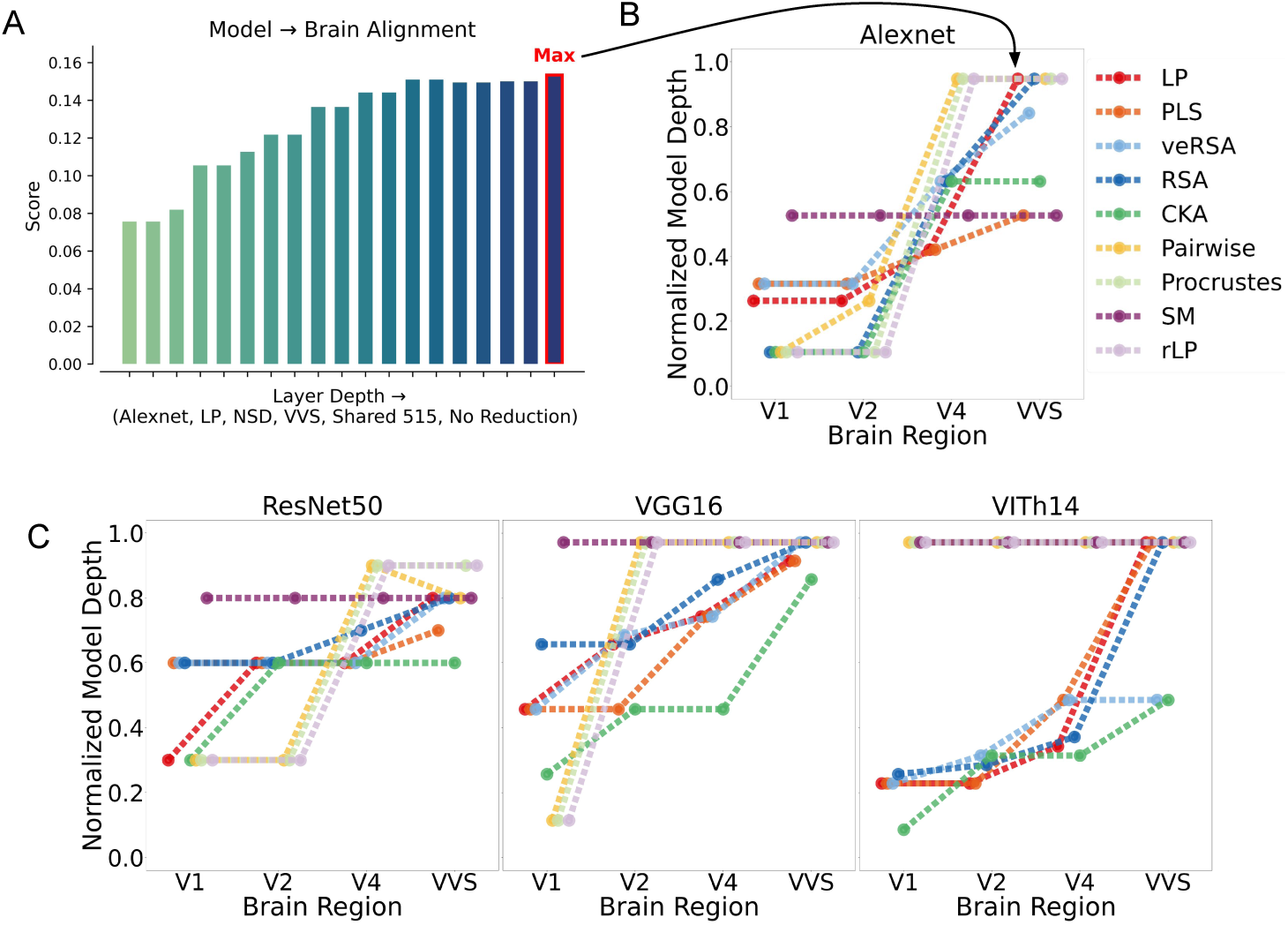
Across measures, higher brain areas tend to best correspond to higher levels in the neural networks. (A) We first find the depth of the best fitting layer for each brain region, identified using the shared 1,000 NSD images. (B) This data is plotted for Alexnet for multiple measures. (C) We repeat this for 3 other standard ImageNet trained models using sparse random projection as dimensionality reduction. Layer depth is normalized with 0 being the first layer and 1 the output layer. A slight jitter is added to the X-axis to make overlapping points more visible.

While this could be evidence for only using the measures that are able to recover this correspondence, we find that multiple of the measures that are able to recover this correspondence between artificial and biological neural systems fail when applied between biological neural systems. Specifically, when the same hierarchy analysis is applied between subjects within NSD, by holding out data from a single biological neural system to replace the artificial one, linear predictivity and several other directional measures fail to map the systems’ V2 and V4 to the corresponding V2 and V4 of the other subjects in the dataset (Fig. A.1). We believe this highlights a fundamental problem for predictive metrics (further highlighted in sections 3.4 and 3.6 and by Avitan & Golan 2026 [54]) where an imbalance in coverage leads to inflated scores, even when the underlying similarity is low. In this specific example, the representations for V1 and VVS are significantly larger than V2 and V4, and while larger representations don’t always mean the representation has larger coverage, the mismatch we see in the brain-to-brain comparisons suggests this is the case.

These failures in a brain-to-brain setting highlight that the choice of measure determines which model layer is said to correspond to which brain area. Because different layers in these networks encode qualitatively different features, layer-to-area mappings are one of the main ways comparative analyses support claims about the kind of computation a brain region performs. Two measures that produce incompatible mappings on the same data cannot both support a correct biological interpretation. We return to what this implies for measure selection in the Discussion.

### 3.2 Measures Exhibit Weak Empirical Correlations

While hierarchical correspondence provides us with a rough idea about the differences between these metrics, it is a coarse measure that can be sensitive to noise when many layers in the artificial neural system have similar scores to the same layers in the biological systems. Therefore a more robust comparison considers how well every layer is mapped rather than just a single one. To do this, we obtain a score from every layer of a network and compare how these scores change depending on the measure. If the hierarchical correspondence problem is due to noise, the correlations of the layer by layer scores across measures should still remain high.

We investigate these between-metric correlations across layers on the network that started the DL revolution, AlexNet ([42]). To do this, we compute similarity scores for every layer across the network, repeating this for all 9 scores, with and without dimensionality reduction, for many datasets (including subsets within a dataset, such as brain region or imageset). We find that the measures are quite distinct, often having small or even negative correlations. We see this within a specific subset of a dataset (Fig. 3 A,B where we only look at relationships in the V1 subregion of NSD, using the shared 1000 image subset, without dimensionality reduction). And continue to see this when collapsing across a variety of datasets (Fig. 3 C).

**Fig. 3.**
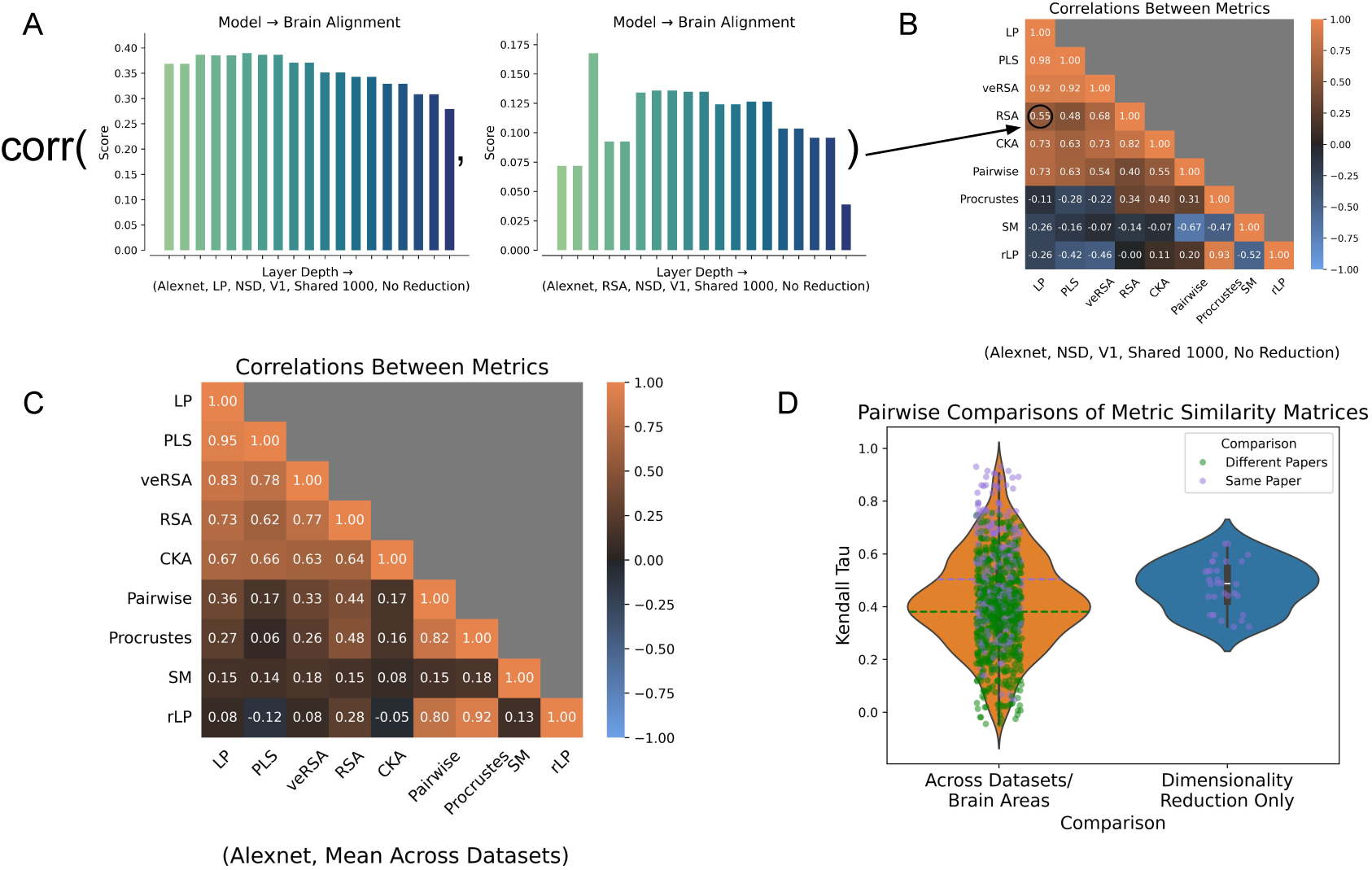
Similarity measures show weak correlations across layers of AlexNet. (A) For each pair of measures, we compute the correlation between their layer-wise similarity scores across AlexNet layers for a given brain region and dataset. (B) This generates a 9×9 Measure Similarity Matrix (MSM) showing how consistently different measures respond across layers. (C) We compute MSMs for all 32 brain region/dataset combinations and average them to reveal overall measure relationships. (D) Pairwise comparisons of MSMs using Kendall’s *τ* show that measures are more consistent within the same dataset than across different datasets. This also highlights, that the use of dimensionality reduction significantly affects the relationships between metrics.

Not only the measure, but other methodological choices in this comparison also matter. For example, many comparison procedures start with a dimensionality reduction such as sparse random projection (SRP) step to effectively deal with high-dimensional representations. While this is often a necessary step to make the analysis computationally feasible, if it affects the results, feasibility is not a sufficient justification. Indeed, we find that the nature of dimensionality reduction (SRP) has major impact on the relationship between metrics. We find that the metric relationships change significantly once SRP is applied (Mean Kendall’s *τ* 0.479, Fig. 3 D, right). This means that dimensionality reduction techniques must also be linked to the scientific question to ensure consistent scientific conclusions.

Another concern is how dataset dependent these results are. The datasets used are used for their size and availability with a preference for more natural images. This means that the dataset is not directly linked to the scientific question and for principled conclusions, varying the datasets should not significantly change the results. We find again that the dataset has a major impact on the relationships between the metrics (Mean Kendall’s *τ* 0.417, Fig. 3 D, left). We see that the relationship of metrics within the same dataset, but across brain regions (Mean Kendall’s *τ* 0.507) is significantly different than the relationship across datasets (Mean Kendall’s *τ* 0.382) and find that the dataset has a greater impact than the depth of the brain area recorded (*p <* 10^−5^). We see that multiple factors affect the relationships between measures including but not limited to dimensionality reduction, the dataset, and the depth of the recordings.

The similarity matrices are not unstructured. Two broad clusters emerge: on one side, more permissive measures such as linear predictivity and RSA, and on the other, stricter measures such as pairwise matching and soft matching. Some measures sit unexpectedly; for instance, reverse linear predictivity falls with the strict matching group despite the common intuition that forward and reverse linear predictivity should behave similarly (we return to this in *Forward Vs. Reverse Linear Predictivity*). This cluster structure highlights how the different measures are sensitive to different things and may imply that tying a certain cluster to your question, such that the cluster of metrics is the only theoretically justifiable one, may be a correct step forward. However, substantially more theoretical work is needed before those recommendations can be concrete, especially since there does not seem to be a clear theoretical separation between the clusters.

### 3.3 Different Measures Yield Different Rankings of Models

While the low correlation across layers of the model when using different metrics is concerning, typically the actual scientific conclusions are made from rankings between models. Each of these models embodies a different hypothesis about neural computation and how well they align with neural data compared to other models determines which hypothesis is correct. For the conclusions drawn from this method to be robust, the rankings of models should not depend strongly on the measure used. We find that they do. Across 25 models, rankings vary dramatically across measures: a model that is ranked at the bottom for one measure can appear close to the top for another (Fig. 4 B). This is not isolated to individual cases, summarizing the rank difference with pairwise rank correlations finds that metrics make very different rankings for models in multiple brain areas. The worst is when comparing to V1 where metrics are even anticorrelated (mean Spearman correlation 0.13, Fig. 4 C) and still low in VVS (mean 0.43, Fig. 4 D), with a similar cluster structure to that seen in Fig. 3.

**Fig. 4.**
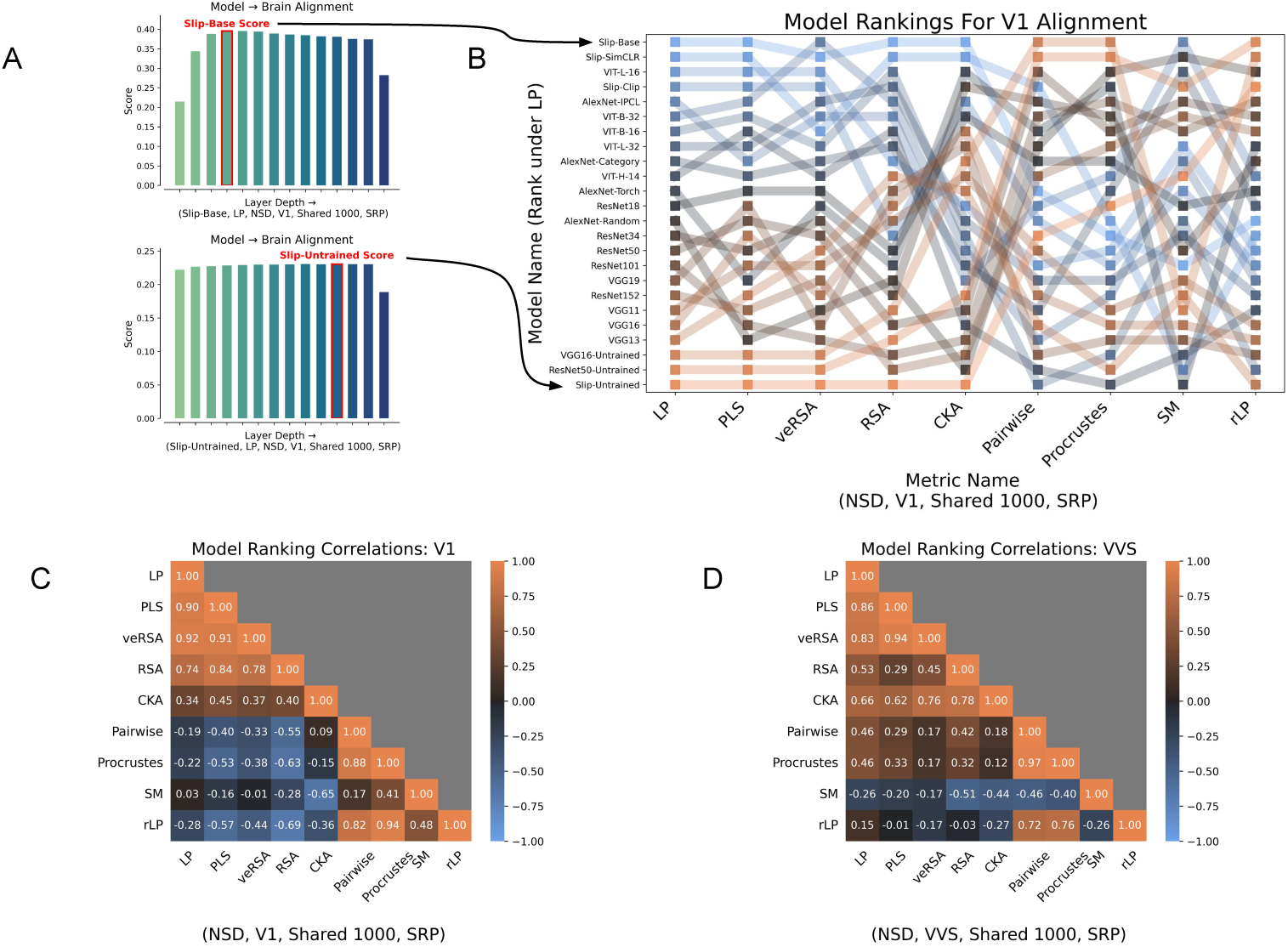
The choice of Measure affects the preferred models. (A) We obtain a score for each model (for a specific region and measure) by first computing similarity with each block of the model and choosing the best one. (B) Using these scores, we rank the models for each measure separately, with the transparent lines indicating how each model’s rankings change from one measure to the next. Models are colored with their rank under linear predictivity, this coloring highlights that some of the best models under linear predictivity end up as the worst under other measures. (C) Similarity matrices of pairwise rank (Spearman) correlations for ranks of models under different measures for V1 (D) and again for VVS.

This means that two studies using the same models and the same dataset of brain data can reach different conclusions about which architectures or training regimes produce the most brain-like networks, solely because of the measure they chose. The ranking is not solely due to a property of the biology, instead it is a property of the measurement.

### 3.4 Forward Vs. Reverse Linear Predictivity

Forward (encoding) and reverse (decoding) predictivity measures are fundamentally different analyses when utilizing the typical approach of investigating stimulus properties (i.e. encoding models to predict stimulus type). In the case of comparing neural systems we see a different relationship where both become equally important. Claims that a model is “brain-like” [28] or that a model’s computations are algorithmically similar to those of the brain [22] implicitly assume representational equivalence between model and brain. Under that assumption, the relationship between artificial and biological neural systems should be symmetric, more specifically, forward and reverse predictivity should show similar results. We instead find low rank correlation of models between forward and reverse predictivity (.15, Fig. 4) and find that the two lead to different conclusions (Fig. 7).

While there are potential issues in treating forward and reverse predictivity as the same, the specific methods we use mitigate this. One such issue is that data from the model and the brain differ significantly in dimensionality. However in many comparisons with VVS both the artificial and biological neural system have a dimensionality of around 10^3^ and we still see low correlations between the two (Fig. 4, D). The other major issue is that recording methods in the biological neural systems can be sparse (electrophysiology) or degraded (fMRI). We mitigate this with our comparisons that utilize dimensionality reduction which degrades the data on the model side and we still see low correlations (Fig. 4, C). The asymmetries that remain imply one of two things: either forward and reverse predictivity capture distinct computational relationships that do not lead to principled scientific conclusions when comparing models to brains, or the representations being compared are not equivalent in the first place — which is a substantive blow to claims of representational similarity.

### 3.5 How Much Alignment Have We Actually Established?

The field typically interprets correspondence between artificial and biological neural networks by comparing it to the correspondence between multiple biological neural systems (taken as a maximum reference point) with an untrained-artificial system providing a baseline. Because untrained models are not always worse than trained models in practice (see fig. 4 and again later in fig. 7), we additionally compute two alternative baselines: the raw pixel values of the image, and a normalized vector of category information. The results depend heavily on the measure. For some measures, the average model sits well above baseline and approaches the correspondence between multiple biological neural systems. For others, the average model is below the pixel or category baseline. This means that the artificial neural system is less similar to biological neural systems than simply the pixel intensities of the stimulus (Fig. 5). In this case the algorithms of the artificial neural network are not only different from humans, but transform the input in a way that explicitly removes the information used in the biological algorithm.

**Fig. 5.**
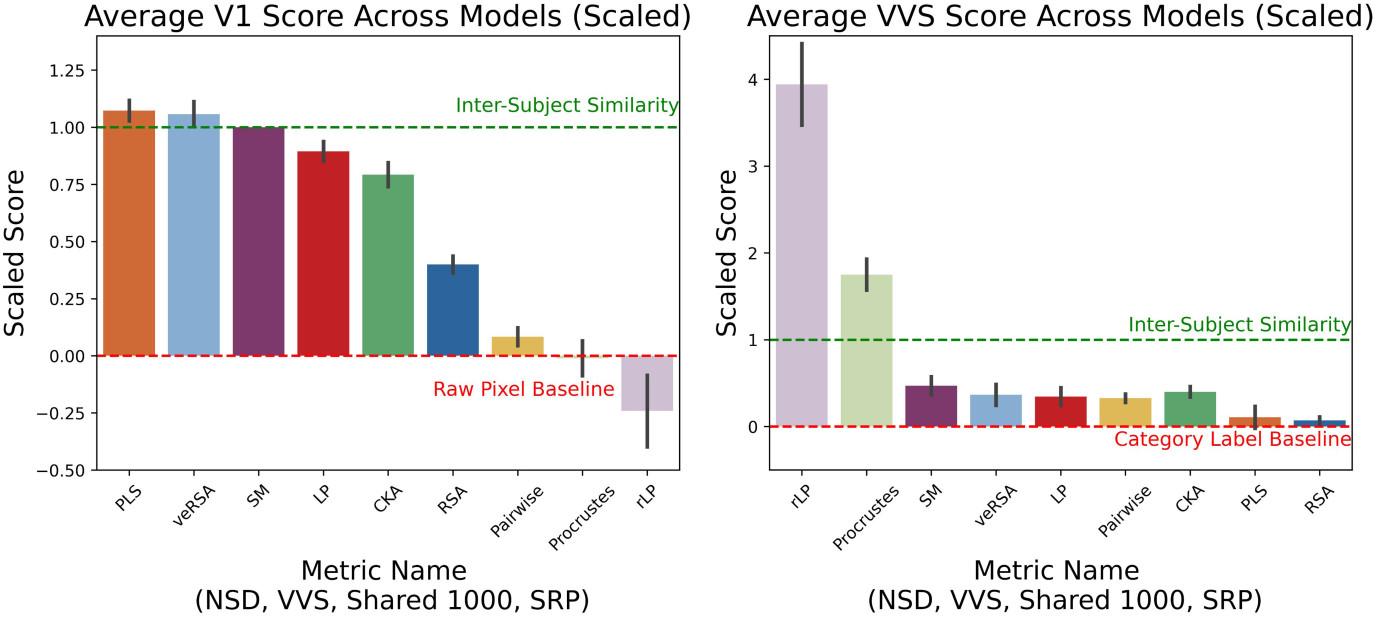
Some measures seem misleadingly good while others appear disappointingly bad. The average similarity score (NSD Shared 1000, with sparse random projection) across many models for V1 (Left) and VVS (Right). The value is scaled by the inter-subject reference point at 1 and either the pixel values for the image (Left) or the category information (Right) at 0. Error bars show *±* standard error.

**Table 1.** Information for the measures utilized in our analysis.

| Metric | Shorthand | Fitting Step? | Symmetrical? | Global or Unit Level? |
| --- | --- | --- | --- | --- |
| Linear Predictivity | LP | Y | N | Global |
| Partial Least Squares Regression | PLS | Y | N | Global |
| Voxel Encoded Representational Similarity Analysis | veRSA | Y | N | Global |
| Representational Similarity Analysis | RSA | N | Y | Global |
| Centered Kernel Alignment | CKA | N | Y | Global |
| Pairwise Matching | Pairwise | Y | N | Unit |
| Procrustes Distance | Procrustes | Y | Y | Global |
| Soft Matching Distance | SM | Y | Y | Unit |
| Reverse Linear Predictivity | rLP | Y | N | Global |

One caveat: models are compared to individual brains, so a fitting measure can, in principle, predict a single subject’s activity better than another subject can. This makes the comparison to the inter-subject reference point less of an absolute bound than it is often treated as. But even setting this aside, the fact that measures disagree on whether current models are near this inter-subject reference point or below a pixel baseline is already a serious problem. “How well do our models capture the brain?” does not have one answer given current practice; it has nine.

### 3.6 Simulating Metric Disagreements

One concern with our previous methods is that they do not distinguish between a measure failure and a measurement failure. Even if the measures perfectly agree, if the measurements we make in biological neural systems are extremely noisy it can look like they disagree. With the black box nature of artificial neural systems, it is hard to tell exactly where the failure is coming from. To address this we utilize a small simulated set of neural responses, differing in interpretable systematic ways to show that the disagreements we observe reflect the properties of the measures themselves. We simulate 50-unit neural populations that respond to a simple 1-dimensional stimulus, with each unit described as a simple radial basis function with a preferred tuning and bandwidth. We then randomly sample 5000 1-d stimuli to extract representations from each artificial system. We construct different populations in which Linear Predictivity (the most popular and flexible measure) reports the two representations as perfectly aligned while another measure reports them as different (Fig. 6). Small structural changes — altering the proportions of units with a given selectivity, adding a shared high-frequency component, or changing the bandwidth of selective units — produce large disagreements between measures with no measurement noise present at all.

**Fig. 6.**
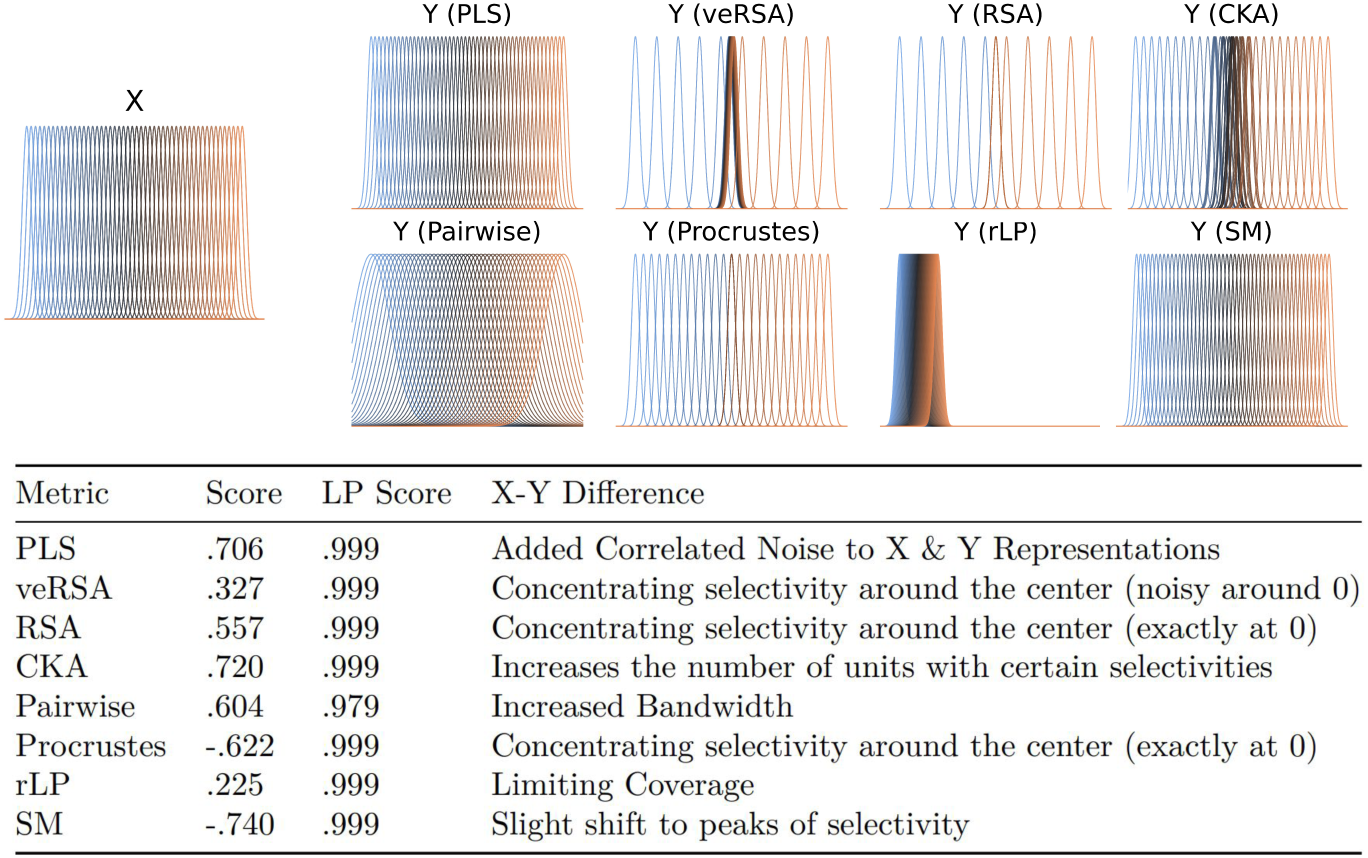
Simulations highlight metric disagreement for functional (non-noise) reasons. Representational Comparisons for two simulated populations of neurons, modeled as radial basis functions 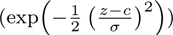 with varying selectivity/center (c) and bandwidth (*σ*). Both X and Y are sets of 50 selective neurons. X is consistent across every comparison. Y varies from X in different ways for each measure, with the goal of having a low score for that measure while keeping a high score for Linear Predictivity. Changes from X to Y are described in the table, exact numbers for each are present in the appendix (Table 2). We sample 5000 “stimuli” uniformly between −10 and 10 to have a 50 by 5000 matrix for the X and Y population that is then compared with Linear Predictivity and the corresponding labeled measure in the same way as done in model-brain (in this case X-Y) comparisons.

**Table 2.** Chosen centers and bandwidths for each representation in Fig 6. 50 centers are chosen, [x,y] means they were chosen equally spaced between x and y, *{*x,y*}* is instead uniformly sampled between x and y, x is all chosen as that value. Percentages indicate what percent of the centers were chosen from the specified set. Bandwidths are constant for all 50 centers. PLS is unique as the selectivity is the same as X, instead correlated noise is added to the representations before calculating the final score.

| Name | X | PLS | veRSA | RSA | CKA |
| --- | --- | --- | --- | --- | --- |
| Center | [-5,5] | [-5,5] | 20% [-5,5], 80% {-0.15,0.15} | 20% [-5,5], 80% 0 | 50% [-5,5], 50% {-1,1} |
| Bandwidth | .2 | .2 | .2 | .2 | .2 |
| Name |  | Pairwise | Procrustes | rLP | SM |
| Center |  | [-5,5] | 50% [-5,5], 50% 0 | [-5,-3] | [-5,5] |
| Bandwidth |  | 1 | .2 | .2 | .2 |

### 3.7 Revisiting Conclusions from Prior Work

While methodological issues are important to investigate, in the end it is only worth significantly upending a subfield when the inferences actually drawn in the literature are incorrect. To see if this is the case we re-examined three prominent results under our nine measures. The three papers were chosen because of data availability and as they served as the first author’s introduction to the field. These were the first and only three papers we examined so any results are not cherry-picked and reflect the state of the subfield. To keep the comparison focused, we only vary the measure used for comparison and do not endorse or criticize other methodological choices (their choice of models, datasets, and so on).

#### 3.7.1 Unsupervised vs. Supervised Learning

One longstanding question in NeuroAI is whether self-supervised networks, arguably more biologically plausible than category-supervised ones, produce better models of the visual cortex [22]. Many self-supervised objectives learn invariant embeddings of different views of the same image, which has a natural analogue in learning stable percepts across eye movements and viewpoint changes. The question remains if these instance level classifications can lead to the same object-level pressures that have long been thought to shape the visual system.

Using the same dataset and models as [22], we obtain conflicting results depending on the measure. Overall, there is a slight bias toward self-supervised learning, consistent with the paper’s conclusion that it is competitive for improving DNN–brain similarity (Fig. 7 C). However, for the paper’s claim to hold, the more basic precondition is that *training* has an effect in the first place — the trained models must outperform their untrained initialization. We scaled the trained scores by subtracting the untrained scores, so any negative value means training lowered the similarity score. Almost a third of the measure/brain-region comparisons indicate that the untrained model outperforms both trained models; half indicate it outperforms at least one trained model. A two-sided paired t-test across subjects per measure finds that several measures show the untrained model significantly beating the trained models, and in one case two of the most commonly used measures — linear predictivity and RSA — yield significant results in *opposite* directions on the same data (Fig. 7 D, note that we caution over-interpreting this result; we discuss this further in the discussion section, “Challenges in standardizing model-brain similarity evaluations”). Our findings suggest that most of the apparent brain-alignment of these models arises from the architecture (most likely the convolutional bias) rather than from the training, and that on this question the choice of measure matters more than the supervised-vs-unsupervised distinction that the original study aimed to investigate.

**Fig. 7.**
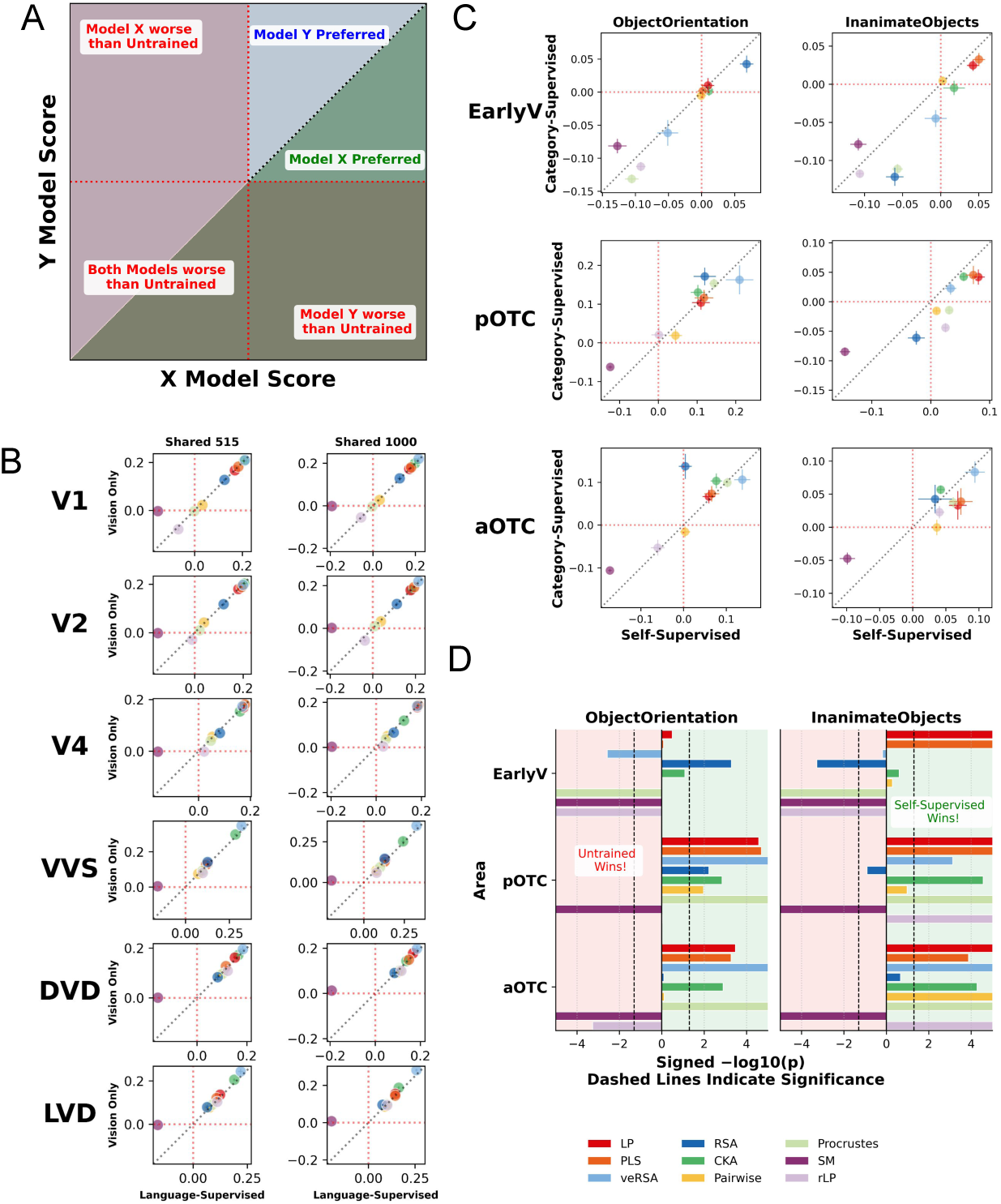
Conclusions about Self-supervised models or multi-modal models fitting brain data better than baseline models are highly fragile to measure choice. We replicate results comparing (C) Unsupervised (IPCL) vs. Category Supervised Learning [22] and (B) Language-Only vs. Language+Vision models [23]. The meaning of each quadrant is defined in (A). We utilize the same datasets as in the papers (ObjectOrientation and InanimateObjects for (C) and NSD for (B)). In both cases we scale the results by the untrained model such that the location of the data point for each metric relative to the dashed lines highlights different potential conclusions. In (C) we see that untrained models do better than both relevant models for many metrics whereas we see in (B) almost all metrics are on-diagonal and no conclusions can be made. (D) Highlights statistical tests comparing the untrained and supervised models as for the conclusions in [22] to hold, training must have an effect. We see statistically significant effects putting either model significantly better than the other depending on the measure. This is the most concerning in early visual cortex (EarlyV) with the Object Orientation dataset showing statistical significance in opposite directions for RSA and LP. The scores for SM are scaled down by a factor of 10 to keep it in the same scale as the rest of the metrics. Error bars show *±* standard error.

#### 3.7.2 The Importance of Language

A second active question is whether networks trained multimodally on visual and text data — arguably a more human-like training signal — align better with the brain, with existing work reaching varying conclusions [23, 24, 33, 55, 56].

Like several of those papers, we use SLIP models, a set of three networks trained on a similar dataset and with similar hyperparameters, differing only in their training objective: language supervision, image-based self-supervision, or both. Almost all of our measures agree that the trained models do better than the untrained model, but between the different training methods there is essentially no consistent effect: most measures sit on the diagonal, and what off-diagonal effects exist are small and unbalanced across measures. Deeper brain areas tilt slightly off-diagonal, but there is no stable preference for either multimodal or single-modality training (Fig. 7 B). Strong claims that multimodal training makes models more brain-like are, on this evidence, not robust: some measures prefer multimodal models, others prefer single-modality models, and the effect sizes are small across the board.

#### 3.7.3 Topographic Networks

A third question is whether matching the topographic structure of the cortex (e.g., the spatial organization of orientation columns) improves alignment with neural activity [28]. Margalit et al. introduced a technique to regulate spatial correlation in trained networks, and it produces pinwheel-like V1 structure reminiscent of earlier work based on independent component analysis [57]. Beyond this appealing result on spatial organization, they made two further claims that depend on a similarity measure: (1) regulating spatial coherence to the level that best matches human cortical topography also yields the best model-brain alignment in ventral temporal cortex (VTC), and (2) their chosen self-supervised objective combined with this level of spatial correlation approaches the inter-subject reference point (referred to as a ceiling in original paper) of alignment in VTC. Together, these claims mean that fitting the right spatial structure produces a network that is “brain-like” both topographically and representationally.

We reanalyzed their setup with minor methodological differences (PLS regression with 25 rather than 1000 components, RSA across images rather than categories, and a different implementation of pairwise matching; see Methods). The measure again has a strong influence on the resulting model-brain alignment scores (Fig. 8).

**Fig. 8.**
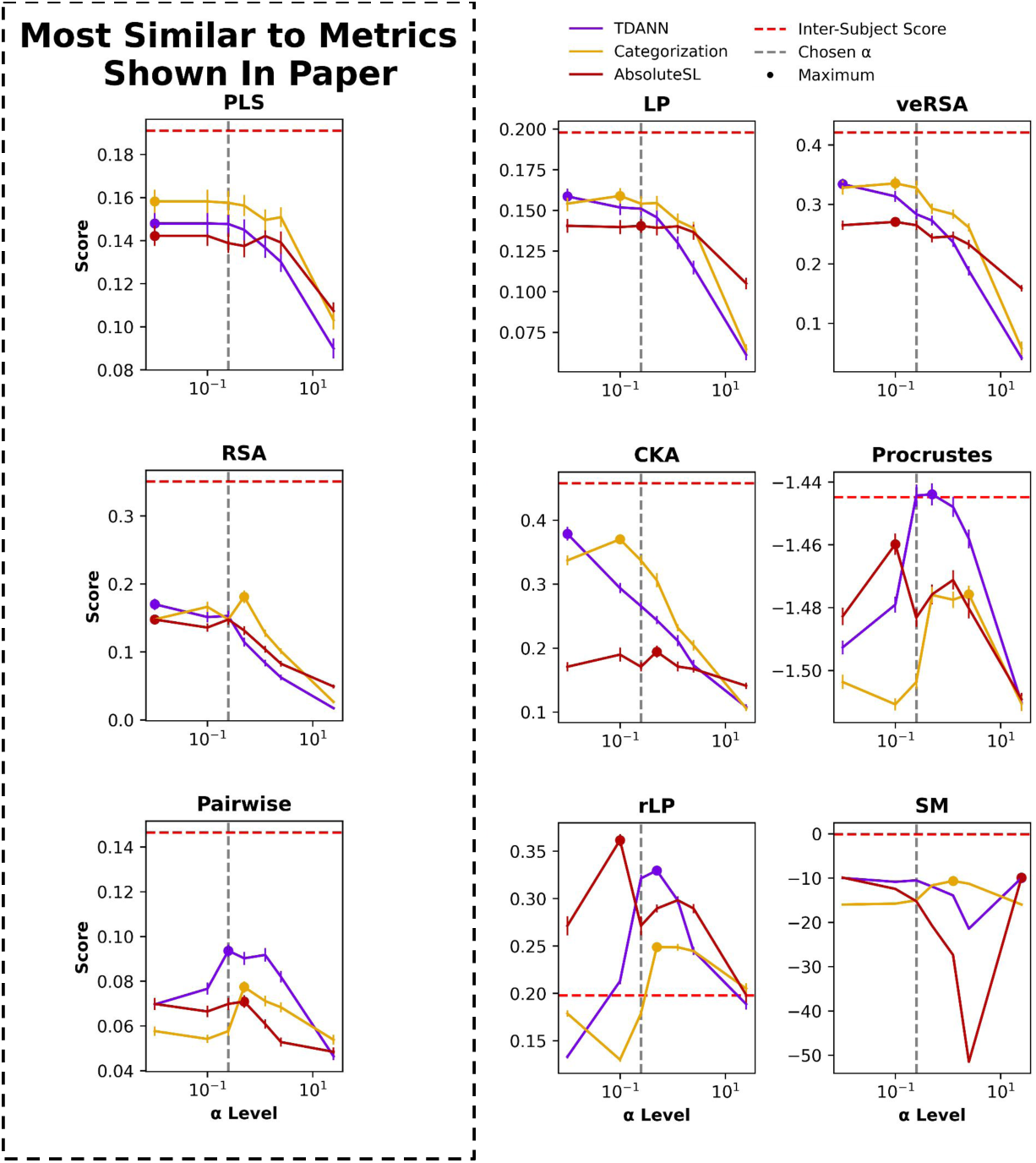
Matching the spatial structure of brains only makes models more brainlike for a limited choice of measures. Replotting Figure 6 (B) from Margalit 2024 [28] with 9 measures (NSD VVS without Dimensionality Reduction). The dots represent which level of spatial correlation leads to the best alignment for each learning rule. The gray line denotes the Paper’s chosen value that matches topographic properties of the brain such as pinwheels. Error bars show *±* standard error.

Only one measure satisfies both (1) the alpha dependence and (2) the training-objective dependence that the paper claims. That measure — pairwise matching — is closest to the measure the paper ultimately selects. The paper does justify this selection on the principled grounds that their PLS regression measure cannot separate the models. That justification is internally consistent, but it is also a textbook instance of the measure-choice problem: when a measure is chosen in part because it separates the hypotheses being tested, the resulting alignment score is no longer independent evidence for the hypothesis. With nine measures, and substantial hyperparameter freedom within each, a determined analyst could produce a post-hoc justification for almost any pattern. The observation that measure choice affects not only the alignment score but the scientific claim that score supports warrants a higher degree of scrutiny on these methods than the field currently applies.

## 4 Discussion

Comparing representations in biological and artificial neural systems has become a dominant analytical approach for understanding the computational structure of brain regions. The ecosystem around this approach has grown rapidly, but much of the methodological discussion has been theoretical rather than empirical [14, 15, 58–60], and there has been little systematic evidence on whether different similarity measures yield similar conclusions when applied to the same data. That gap leaves the choice of measure — which is one of the most consequential methodological decisions in this entire program — governed largely by convention rather than by principle.

Several prior studies have compared similarity measures against specific theoretical or empirical criteria, typically across a small set of classical measures. Han et al. [61] evaluated measures on system identification — their ability to recover a ground-truth model from a set of candidates. Kornblith et al. [32] argued that a good similarity measure should be invariant to orthogonal transformations and isotropic scaling but not to arbitrary invertible linear transformations, and found CKA best satisfied these criteria. Ding et al. [62] examined the sensitivity of CCA, CKA, and orthogonal Procrustes distance to factors that do not affect the functional behavior of models. These studies largely focus on classical measures (RSA, CKA, encoding analysis) and on theoretical properties rather than on downstream scientific conclusions. While there are theoretical reasons to prefer some measures over others, the field has not reached a consensus on when specific measures should be used, so in the interim the breadth of measures in common use should all be taken seriously as candidates.

### Inconsistencies and their implications

Our evaluation across multiple datasets and models shows that the choice of measure causes inconsistencies in alignment scores within and across models, and that these inconsistencies undermine several claims in the NeuroAI literature. The effects of unsupervised learning and of language supervision in producing more brain-like models remain unclear, with different measures yielding different answers and effect sizes that are often small relative to differences from untrained networks (Section 3.7). These instabilities matter because subsequent work has built theoretical positions on top of these comparisons. The reported advantage of contrastive self-supervised models has been generalized into a claim that contrastive coding provides “a unifying account of object category emergence and representation in the human brain” [8], and is cited to justify the task component of the loss in later model-fitting work such as TDANN [28]. The reported advantage of language-aligned over image-only models has been used to argue that “transforming visual input into rich semantic scene descriptions may be a central objective of the visual system” [55] and that language causally shapes the ventral stream [63]. Claims of this scope are claims about the overall similarity between systems, and should rest on evidence that survives the choice of similarity measure. Some of the underlying inconsistencies may attenuate as more diverse models are compared against more brain data, since the marginal effect of any single manipulation may be stable in aggregate even when individual pairwise comparisons are not. But such comprehensive analyses are resource-intensive and unavailable for most individual studies — and would not retroactively justify conclusions already published on the basis of a single measure.

### Common considerations in measure selection

This analysis treats all measures as a priori indistinguishable. However, each measure has different properties and invariance that tie them to specific questions. Studies that aim to provide computational accounts of neural tuning functions, for example, require measures that are sensitive to rotations but invariant to neuron-index permutations, which narrows the pool of appropriate choices. Prior work on inter-model similarity and on representational alignment with the brain [52, 54, 64, 65] identifies other contexts where specific measures are better matched to the scientific question. In those cases, the finding that measures disagree does not undermine the conclusion — as the chosen measure is the only theoretically justified one for the question at hand. But as we discuss in the introduction the broader literature provides reasons for measure choice such as scoping out more permissive measures [28] that do not isolate a single measure. Given the incentives to produce conclusions consistent with the intuitions of the field, this creates a real risk of motivated selection, even if unintentional.

### Challenges in standardizing model-brain similarity evaluations

The hyperparameter space inside these analyses is larger than measure choice alone. The metric used to compare RDMs in RSA varies (Kendall’s *τ*, Spearman, CKA, and others). The Procrustes distance has a tuning parameter *α* that can vary between 0 and 1 (we use *α* = 1, as recommended in the original paper). There is also substantial freedom in which model activations to compare: best-fitting layer, final embedding, or multiple layers combined. A wide hyperparameter space spanning multiple stages of a comparison is exactly the setup in which cherry-picking — intentional or not — is most likely to occur, and it requires that each of these choices be made and documented with care.

This same sensitivity to experimenter choice is why our analysis does not include a deeper statistical treatment. We omit it for two reasons. First, we comment on the eventual scientific conclusions: even if none of our results in Section 3.7 were significant — which Fig. 7 shows is not the case — that non-significance would itself directly support our claims. More importantly, the multi-scale structure of neural data leaves the appropriate statistical method underdetermined, and each procedure rests on choices that changes the numbers they produce. Bootstrapped confidence intervals are one option, but their value depends heavily on how the resampling is defined (across subjects, across splits, across bootstrapped samples of stimuli), and different studies choose differently. Reference points (sometimes referred to as a noise-ceiling) differ similarly across the literature: inter-subject alignment, within-subject consistency across repeated trials, and ideal-observer or pixel-based lower bounds are all in use, rescaling what counts as strong or weak results. Any single treatment we reported would therefore embed one defensible convention among many rather than a settled answer. Future work on model-brain alignment should prioritize standardized and principled statistical practice.

### Reconciling discrepancies

The inconsistency between measures is not purely a flaw to be engineered away. Different measures are sensitive to different aspects of representations — geometry, information content, unit-level correspondence — and the disagreement between them is itself informative about what *kind* of similarity holds between a model and a brain. Claims should emphasize not only the degree of similarity but also the nature of that similarity.

Using multiple measures together can provide complementary insight. Such comparisons also offer an opportunity to revisit and critically analyze the implicit assumptions of existing comparison techniques, and to develop new tools that address their limitations. For instance, the asymmetry we observe between linear predictivity (brain variance explained by models) and reverse linear predictivity (model variance explained by brains) highlights the limits of unidirectional measures and the need to interpret them carefully. As with asymmetries in human similarity judgments, such as in the Korea–China analogy [66], model–brain comparisons can capture broad computational correspondences without guaranteeing bidirectional equivalence. Models may align with neural representations in ways that are useful for prediction while diverging in structure or constraints that affect reverse predictivity. Rather than treating similarity measures as definitive indicators of representational equivalence, they should be interpreted in terms of what aspects of neural computation are recoverable in each direction. We emphasize that reporting multiple measures does not by itself resolve the problem: doing so usefully requires explicit discussion of what each measure treats as similar and what theoretical assumptions it carries. Reporting multiple numbers without that discussion is not meaningfully more informative than reporting one.

### Alternate emerging tools for representational similarity

Several tools not studied here are also available for assessing representational similarity. These include interventional measures that go beyond correlations by asking how substituting one representation for another affects downstream computations [67], measures that do not assume a Euclidean representational space [68], and measures that quantify hierarchical brain-like structure in a network [53]. A thorough comparison of these tools with the measures studied here is left to future work.

### Similar problems in other domains

Recent work suggests the measure-choice problem is not limited to vision. Cloos et al. [69] found similar results for temporal data and RNNs, where Angular Procrustes, linear predictivity, and CKA prioritize different principal components. In the language domain the problem appears more severe: AlKhamissi et al. [70] (their Supplementary Fig. 10) report that the bottom three models under RSA are the top three under linear predictivity — measures with a rank correlation of at least 0.53 in our vision data (Fig. 4) appear anticorrelated for language. Feghhi and Hadidi et al. [71] show that much of the apparent ability of LLMs to predict brain activity is captured by trivial features such as temporal autocorrelation, sentence length, and sentence position. The measure-choice problem is a cross-domain issue, not a vision-specific one.

Reproducibility concerns are also familiar from neighboring fields. Neuroscience and psychology have grappled with similar issues [72–75], and preregistration has been proposed as one partial remedy [76, 77]. Strict preregistration is difficult in NeuroAI, where analytical approaches often need to evolve during a project. We recommend that researchers document methodological assumptions and analytical rationales before implementation — a form of self-preregistration that improves transparency and reproducibility without constraining necessary methodological flexibility.

### Theoretical connections between similarity metrics

Recent work has made significant progress on unifying the fragmented landscape of representational similarity measures by uncovering deep theoretical connections between them. Harvey et al. [78] bridge two broad categories — methods that learn explicit mappings (e.g., shape distances) and methods that rely on summary statistics (e.g., normalized Bures similarity) — by showing that the cosine of the Riemannian shape distance equals normalized Bures similarity. Williams et al. [37] show the equivalence of RSA and CKA when a mean-centering step is added to RSA, simplifying the interpretation of both measures. Harvey et al. [79] connect CKA, CCA, and Procrustes distance to a decoding perspective, showing that they quantify the alignment between optimal linear readouts across tasks. These theoretical results clarify the relationships between measures and provide a unifying framework for interpreting their geometric and informational content. Our empirical results complement that work by characterizing relationships among a broader set of measures whose theoretical connections remain poorly understood.

### The importance of open science in NeuroAI

This analysis was only possible because of open science. Every claim we reanalyze depends on datasets and trained models that the original authors released publicly, and in each case the original authors took care to vary a single hyperparameter at a time, isolating the effect the paper was testing. Two implications follow: continued publication of full data and code should remain standard practice in NeuroAI, and any claim that rests on a similarity score alone — without independent mechanistic, behavioral, or ablation-based evidence — should be revisited as a default, not as an exceptional case.

### Navigating the challenges of an emerging field

NeuroAI is a young field, and the tools for evaluating the alignment between brains and models are still being worked out. Unlike traditional neuroscience, which often works in lower-dimensional feature spaces where representational comparisons can cleanly recover function [80] or identify the correct underlying model [81], model–brain alignment operates in very high-dimensional representational spaces with methodological complexities that are only now becoming clear.

Our findings argue for adapting methodology to match these complexities. Rather than relying on a single similarity score, studies should design questions to be linked to the properties of a single or even multiple measures and interpret their agreement and disagreement explicitly. More rigorous evaluation frameworks — on top of the ones that tie the choice of measure to the specific scientific question being asked — such as ones that thoroughly evaluate how methodological choices affect the results are needed if claims about model-brain alignment are to function as reliable evidence about the brain.

## 5 Code Availability

A GitHub repository with the data and code to reproduce all figures and experiments is provided here: https://github.com/anshksoni/NeuroAIMetrics

## 6 Data Availability

Data produced from the analysis in this paper can be found in the GitHub repository at https://github.com/anshksoni/NeuroAIMetrics. All data required to reproduce the analysis come from publicly accessible datasets, which are listed in the repository’s README.

## 7 Author Contributions

AS conceptualized the study, developed the methodology, set up the data, ran the analysis, and wrote the paper. SS and MM conceptualized the study and edited the paper. KK and MK conceptualized the study, developed the methodology, wrote and edited the paper, and supervised the work. Funding was acquired by AS (NSF GRFP), KK, and MK.

## 8 Funding Acknowledgment

This material is based upon work supported by the National Science Foundation Graduate Research Fellowship under Grant No. DGE2236662.

---

**Fig. A.1.**
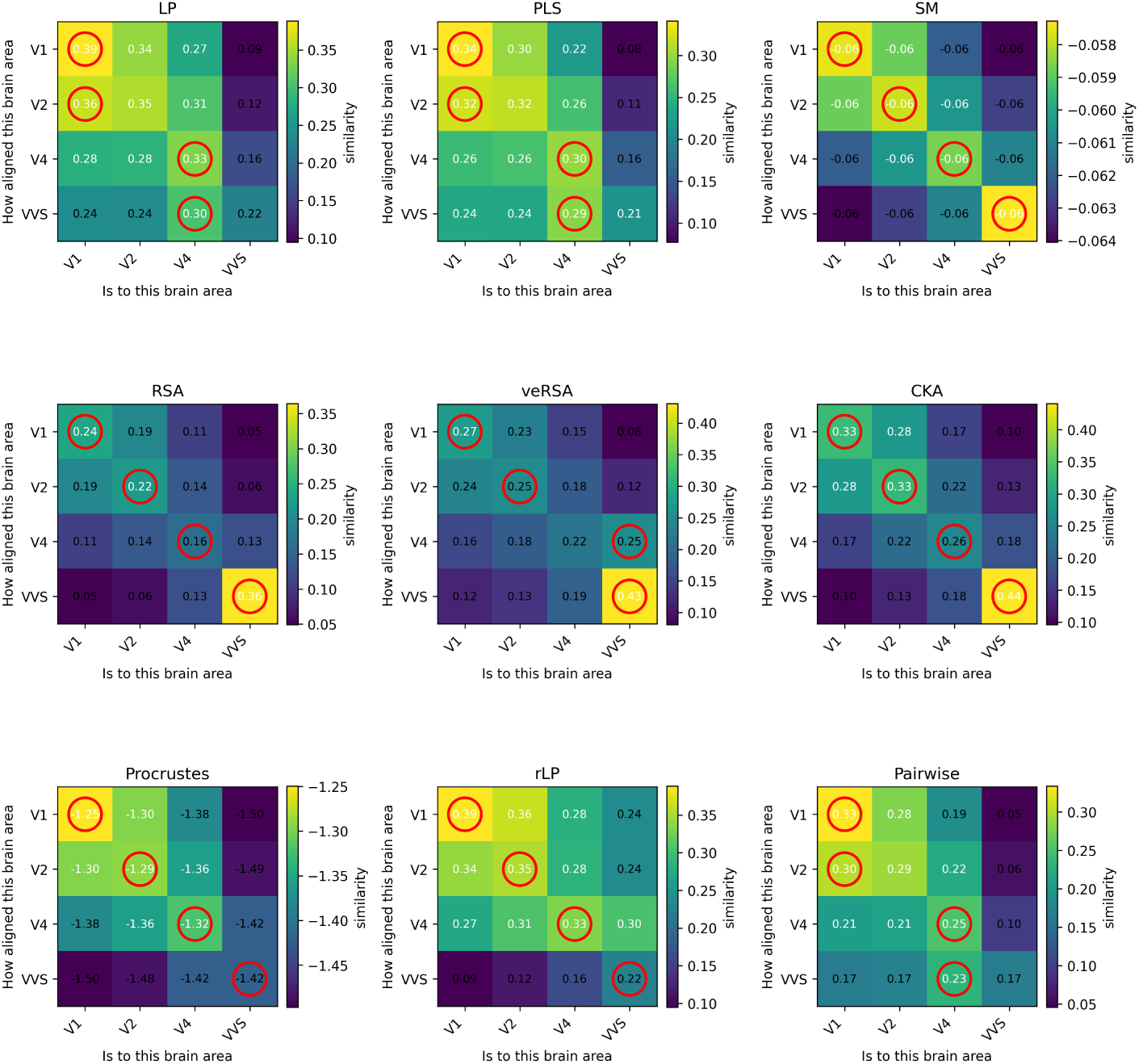
Hierarchy is not recovered when utilizing the model-to-brain method on brain-to-brain. Hierarchy comparisons between subjects in NSD shared 1000. We align each brain area to the other brain areas across subjects, the Y axis indicated the source and the X the target in every metric, therefore the asymmetrical measures have different upper and lower triangles. rLP is therefore also the transpose of LP. Akin to model-brain comparisons, we only compare hierarchy across subjects (e.g. subject one’s V1 is never aligned to their V2) We see that metrics like LP are unable to recover hierarchy (indicated by the off-diagonal red circles).

